# Global Phylogeography and Microdiversity of the Marine Diazotrophic Cyanobacteria *Trichodesmium* and UCYN-A

**DOI:** 10.1101/2024.07.12.603225

**Authors:** Angie Nguyen, Lucas J. Ustick, Alyse A. Larkin, Adam C. Martiny

## Abstract

Cyanobacterial diazotrophs, specifically the genera *Trichodesmium* and UCYN-A, play a pivotal role in marine nitrogen cycling through their capacity for nitrogen fixation. Despite their global distribution, the microdiversity and environmental drivers of these diazotrophs remain underexplored. This study provides a comprehensive analysis of the global diversity and distribution of *Trichodesmium* and UCYN-A using the nitrogenase gene (*nifH*) as a genetic marker. We sequenced 954 samples from the Pacific, Atlantic, and Indian Oceans as part of the Bio-GO-SHIP project. Our results reveal significant phylogenetic and biogeographic differences between and within the two genera. *Trichodesmium* exhibited greater microdiversity compared to UCYN-A, with clades showing region-specific distribution. *Trichodesmium* clades were primarily influenced by temperature and nutrient availability, and particularly frequent in regions of phosphorus stress. In contrast UCYN-A was found in regions of iron stress. UCYN-A clades demonstrated a more homogeneous distributions, with a single sequencing variant within the UCYN-A1 clade dominating across varied environments. The biogeographic patterns and environmental correlations of *Trichodesmium* and UCYN-A highlight the role of microdiversity in their ecological adaptation and reflect their different ecological strategies. This study underscores the importance of characterizing the global patterns of fine-scale genetic diversity to better understand the functional roles and distribution of marine nitrogen-fixing cyanobacteria.

## Introduction

Microbial biogeography and phylogenetic diversity are intricately linked, with genetic variations as small as 0.5% translating into significant physiological differences among microorganisms (Fuhrman and Campbell 1998). Such genetic differences, referred to as “microdiversity,” can have profound implications for microbial adaptation and ecological functions. In many bacterial species, microdiversity leads to the formation of distinct clades, which are genetically cohesive populations adapted to specific environmental conditions (Cohan 2002). The concept of ecologically distinct clades is well illustrated in the cyanobacterium *Prochlorococcus*, which is divided into multiple clades that are adapted to varying light, temperature, and nutrient conditions in marine environments (Johnson et al. 2006; Martiny et al. 2009; Rusch et al. 2010). This study extends the exploration of microbial microdiversity to cyanobacterial diazotrophs, focusing on the globally significant genera *Trichodesmium* and *Candidatus Atelocyanobacterium thalassa* (UCYN-A), both of which contribute to marine nutrient cycling by fixing atmospheric nitrogen into bioavailable forms (Zehr 2011).

*Trichodesmium*, a free-living cyanobacterium, is known for its ability to form large colonies and bloom in nutrient-poor, warm oceanic regions, particularly in the Atlantic Ocean and places where temperatures exceed 25°C (Davis and McGillicuddy 2006; Dugdale, Menzel, and Ryther 1961; Gradoville et al. 2017; Pierella Karlusich et al. 2021; Rouco et al. 2014; Subramaniam et al. 2001). Previous phylogenetic analyses have divided *Trichodesmium* into four clades (Tricho I, II, III, and IV), with Clades I and III being the most well-documented based on the nitrogenase (nifH) gene(Hynes et al. 2012; Orcutt et al. 2002). Clade II, in particular, remains poorly understood due to a lack of comprehensive reference sequences (Lundgren et al. 2005). While the isolate IMS101 from clade III has been used in laboratory experiments, its applicability to Earth system models has been questioned, as regional studies have shown this clade to be less abundant than others in the environment (Gradoville et al. 2014; Hmelo, Van Mooy, and Mincer 2012; Rouco et al. 2014, 2018; Rouco, Haley, and Dyhrman 2016). Despite extensive studies on *Trichodesmium* at the genus level, there is a need for a deeper understanding of the biogeographic distribution and microdiversity of its clades to elucidate their ecological roles and adaptations.

Similarly, *Candidatus Atelocyanobacterium thalassa* or UCYN-A is a widespread marine diazotroph that forms symbiotic relationships with eukaryotic hosts, facilitating its role in nitrogen fixation across diverse marine environments (Martínez-Pérez et al. 2016; Zehr 2011). Phylogenetic analyses of UCYN-A, based on the *nifH* gene, have identified multiple clades (UCYN-A1 to -A6), each exhibiting distinct geographical distributions and environmental associations (Cornejo-Castillo et al. 2019; Farnelid et al. 2016; Turk-Kubo et al. 2017). However, it is still unclear what are the exact drivers of UCYN-A abundance as a genus and how widespread each clade is globally. UCYN-A has become an increasingly significant model organism as it was recently shown to be tightly integrated into its host algal cell, suggesting it is an early evolutionary stage nitrogen fixing organelle termed a “nitroplast” (Coale et al. 2024). This highlights the necessity for comprehensive sequencing and analysis to better understand the distribution, diversity, and ecological significance of UCYN-A clades in the global ocean.

To address these gaps in knowledge, we conducted a global biogeographic analysis of the phylogeography and microdiversity of *Trichodesmium* and UCYN-A using *nifH* gene sequencing across over 1,000 seawater samples from the Pacific, Atlantic, and Indian Oceans, as part of the Bio-GO-SHIP project. By characterizing the phylogenetic diversity and mapping the global distributions of these cyanobacterial diazotrophs, we aim to uncover the environmental drivers and functional differences that shape their ecological niches. This study provides a detailed analysis of the biogeographic patterns and microdiversity of *Trichodesmium* and UCYN-A, contributing to our understanding of their roles in marine ecosystems and informing future research on the adaptation and evolution of marine microorganisms.

## Methods

### Seawater Collection

Between 2-10 L of surface water was collected from each cruise using the Niskin rosette system or the ship’s circulating seawater system. Samples were filtered through a 0.22 µm pore size Sterivex filter (Millipore, Darmstadt, Germany) and preserved with a lysis buffer and stored at −20°C before extraction.

### DNA Extraction

180 mL lysozyme buffer (50 mg/L) was added to each Sterivex containing seawater before incubation at 37°C for 30 min. Then, 180 mL proteinase K (1 mg/mL) and 100 mL sodium dodecyl sulfate was added to the filter before overnight incubation at 55°C on a plate shaker. The filtered liquid, along with sodium acetate and cold isopropanol, was divided into microcentrifuge tubes and then incubated for >2 h at -20°C. The samples were then centrifuged at 15,000 g at 4°C for 30 min. A DNA pellet was isolated by disposing of the supernatant liquid. The pellet was then suspended in TE buffer for 30 min in a 37°C water bath. Following that, the DNA was extracted using a genomic DNA Clean and Concentrator kit (Zymo Corp., Irvine, CA, USA).

### PCR

The concentration of the extracted DNA samples was measured using a Qubit dsDNA HS assay kit (Life Technologies, Carlsbad, CA, USA). The samples were then diluted to 2 µg/mL with TE buffer. The 2 µg/mL samples were then amplified using a nested polymerase chain reaction (nested PCR) optimized for *nifH* according to the (Zehr, Mellon, and Zani 1998) protocol. Modifications to the original protocol were based on tests using environmental samples and a gradient of possible reaction conditions. PCR reactions are done on a C1000 Bio-Rad Thermocycler with 20uL of PCR mixture including 10uL AccuStart II PCR Toughmix (QuantaBio, Beverly, MA), 4uL MilliQ, 1 μL each of forward and reverse primers, and 4uL diluted DNA. For the primary PCR the reverse primers are nifH3 and nifH4 and the DNA template is the extracted DNA. The C1000 Touch Thermo Cycler (Bio-Rad) reaction conditions are 94°C for 1 min, 47°C for 1:30 min, and 72°C for 1:30 min repeated for 30 cycles with a final extension at 72 °C for 10 min before holding at 4 °C. For the secondary PCR the reverse primers are nifH1 and nifH2, and the DNA template is the amplified DNA from the primary PCR. The reaction conditions are 94°C for 1 min, 54°C for 1:30 min, and 72°C for 1:30 min repeated for 30 cycles with a final extension at 72 °C for 10 min before holding at 4 °C. The nested PCR products were then cleaned using a Zymo 96 well Clean and Concentrator Kit (Zymo Corp., Irvine, CA, USA).

### nifH Amplicon Sequencing

Illumina specific transposase adapters (i5 and i7) were added to the *nifH* amplicons. The amplicons were mixed with 1 uL of each adaptor as well as PCR mix and run under the following conditions: 10 cycles of 94°C for 1min, 54°C for 1 min, and 72°C for 1:30 min, and a final extension step at 72 °C for 10 min. The concentration of each sample was determined using fluorescence via a gel and a standardized curve of samples of known concentrations (determined using a Qubit dsDNA HS assay kit (Life Technologies, Carlsbad, CA, USA)). After pooling the samples, large fragments were removed using Agencourt AMPure XP beads (Beckman Coulter Inc., Brea, CA). Finally, a Bioanalyzer chip (Agilent, Santa Clara, CA) was used to check the size distribution of the pool before sequencing it at the UCI Genomics Center.

### Data Analysis

The QIIME 2 pipeline (Bolyen et al. 2019) was used to demultiplex, trim, and quality filter the nifH amplicon sequence data. The representative sequences were then translated into amino acid sequences via the EMBOSS Transeq command line (Rice, Longden, and Bleasby 2000). Ensembl BLAST (Camacho et al. 2009) was used to align protein sequences to a reference database of known *nifH* sequences to identify *Trichodesmium* and UCYN-A sequences.

### Sequence Analysis

Across all samples, the 100 most abundant amplicon sequence variants (ASVs) were identified for both *Trichodesmium* and UCYN-A. The top 100 ASVs were combined with curated sequences to contextualize our resulting trees. For *Trichodesmium* we included *nifH* sequences from *Trichodesmium thiebautii* H9-4 (LAMW01000105.1) and *Trichodesmium* sp. IMS101 (AF167538.1) (Dominic et al. 2000; Walworth et al. 2015). For UCYN-A we included *nifH* sequences from NC_013771.1 (Tripp et al. 2010), AF299420.1 (Zehr et al. 2001), amplicon sequences from (Cornejo-Castillo et al. 2019), and consensus sequences from (Turk-Kubo et al. 2017). We aligned and trimmed these reads using muscle 5.1 (Edgar 2004) and trimAL 1.4.1 with the -gappyout argument (Capella-Gutiérrez, Silla-Martínez, and Gabaldón 2009). The best tree model was determined using megaX (Stecher, Tamura, and Kumar 2020). Based on this analysis a GTRGAMMA model was selected for both trees. Phylogenetic trees were created using raxml-8.2.12 with the raxmlHPC-PTHREADS-SSE3 command (Stamatakis 2014) with AY768418.1 as an outgroup. Clade annotations were applied based on the resulting trees. Trees were explored using the ggtree package in R (R Core Team 2022; Yu et al. 2017).

### Statistical Analysis

All statistical analysis was done using R version 4.2.1 (R Core Team 2022). Shannon diversity was calculated on the relative abundance of all ASVs using the “diversity” function from the vegan package version 2.6-2 (Oksanen et al. 2022). Bray-curtis distances were calculated on the relative abundances of all ASVs using the “vegdist” function and resulting PCOA decomposition was done using the “wcmdscale” function from the vegan package (Oksanen et al. 2022). Correlations between environmental parameters and the PCOA were calculated using “envfit” from the vegan package (Oksanen et al. 2022). Spearman correlations were calculated on the relative abundances of *Trichodesmium* and UCYN-A along with each clade using the cor.test function from the base R stats package (R Core Team 2022). Nutrient stress values were obtained from (Ustick et al. 2021). These measurements have previously been linked to surface nutrient concentrations and proposed as an indicator of nutrient limitation (Garcia et al. 2020; Martiny et al. 2020; Ustick et al. 2021). We used Ω P high, Ω Fe medium, and Ω N high, which are derived from the following genes: alkaline phosphatase genes (phoA, phoX) (Ω P), Fe uptake transporter genes (cirA, expD, febB, fepB/C, tolQ, tonB) (Ω Fe), and nitrite and nitrate assimilation and uptake genes (focA, moaA-E, moeA, napA, narB, nirA) (Ω N).

### Figure Generation

Phylogenetic trees were visualized using ITOLv6 (Letunic and Bork 2024). All non-tree figures were created using ggplot2 in R and arranged using the ggpubr packages (Kassambara 2020; Wickham 2016). All maps were made using ggplot2 and rnaturalearthdata packages (South 2017).

### Code and Data Availability

All code used in this analysis can be found at https://zenodo.org/doi/10.5281/zenodo.12636629. ASV sequences, ASV counts, and associated metadata is included in supplemental tables 1-4.

## Results

In order to capture the global distributions and environmental drivers of *Trichodesmium* and UCYN-A we collected 954 surface ocean water samples as part of Bio-GO-SHIP and sequences the nitrogenase gene *nifH* (Figure 1). We combined this novel data set with existing studies resulting in a collection of 1102 samples (Gradoville et al. 2017; Raes et al. 2020). We characterized the phylogeny of the top 100 most abundant sequences and grouped them into existing known major clades of *Trichodesmium* and UCYN-A. We then explored the global distributions of our sequences and linked these distributions to environmental measurements to determine the niche space of each clade.

**Figure 1:**
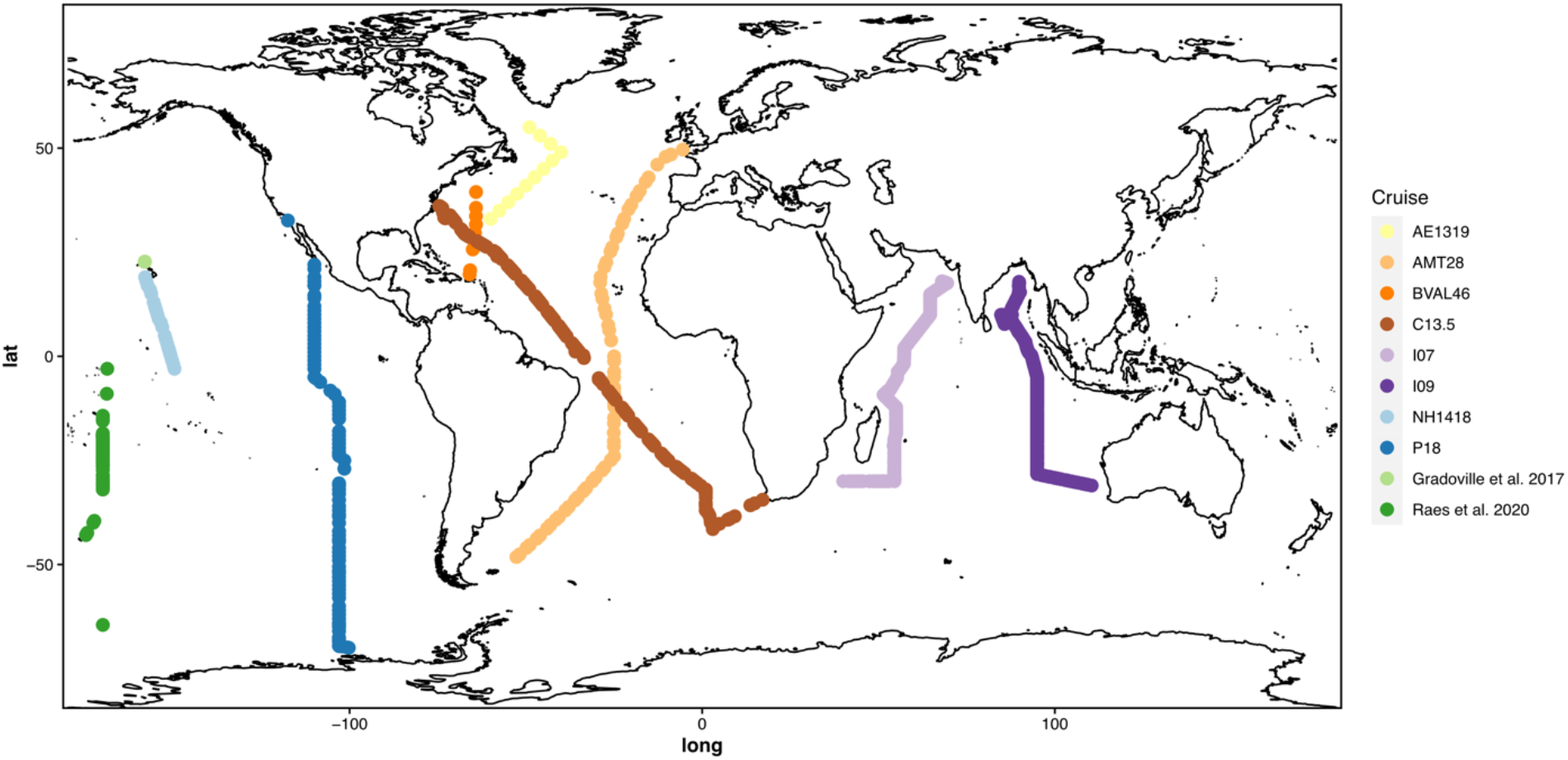
Global distribution of samples.

Phylogenetic analysis of the *nifH* sequences captured known diversity as well as introducing novel representatives. We identified the 100 most abundant ASVs across all our samples. These ASVs constituted 98% of total *Trichodesmium* sequences and 94% of total UCYN-A sequences, thus, we focused on characterizing these sequences further. We created phylogenetic trees for our most abundant sequences and contextualized them using known *nifH* sequences (Figure 2A, B). *Trichodesmium* has previously characterized *nifH* sequences for only two of its four clades. Our tree reconstructed four distinct clades of *Trichodesmium*, which were present in 60%-80% of our tree bootstraps (Figure 2A). Because two of the clades have no previously characterized *nifH* sequences, we inferred the clades based on their proximity to clade I and III. We named these clades UII and UIV since they appear to match clades II and IV respectively. Of the top 100 most abundant ASVs corresponding to *Trichodesmium*, 75 corresponded to the clade Tricho I, 11 to Tricho UII, 6 to Tricho III, and 8 to Tricho UIV. UCYN-A has previously characterized sequence representatives for each of its clades, and our tree captured the four major clades with high confidence (82-95% of bootstraps). Of the top 100 most abundant ASVs corresponding to UCYN-A, 72 corresponded to UCYN-A1, 4 to UCYN-A2, 16 to UCYN-A3, 1 to UCYN-A4, 5 to UCYN-A5, 1 to UCYN-A6 and 1 did not fall within any known clade. Of the top 100 ASVs from *Trichodesmium* and UCYN-A, only a few ASVs made up the majority of acquired sequences (Figure 2C, D). The most abundant *Trichodesmium* ASV, from Tricho I, made up 35% of sequences alone. Although many ASVs were associated with Tricho I, as seen in Figure 2A, only two ASVs were highly abundant (Figure 2C). The second most abundant ASV in *Trichodesmium* was part of clade Tricho UII and made up 26% of total sequences. The rest of the ASVs within the top 100 most abundant represented <5% of total sequences each. Similarly, one ASV within UCYN-A1 made up 45% of all UCYN-A sequences. The second most abundant ASV fell within UCYN-A3 (13%), the third within UCYN-A2 (12%), and the fourth within UCYN-A3 (7.2%). Afterwards, there was a sharp decline in abundance of the remaining ASVs (<2% each). Clades UCYN-A4, UCYN-A5, UCYN-A6, and ASV 50 (which did not fall within a stable clade) combined only represented 0.54% of total sequences and thus were excluded from further analysis. When comparing the alpha diversity of *Trichodesmium* and UCYN-A, we observed a significantly higher average Shannon index for *Trichodesmium* than UCYN-A (*Trichodesmium* mean = 0.55, UCYN-A mean = 0.30, t-test, p < 2e-16)(Figure 2E). In sum, both *Trichodesmium* and UCYN-A demonstrated highly skewed abundance distributions with only 3-4 ASVs dominating across the global surface ocean.

**Figure 2:**
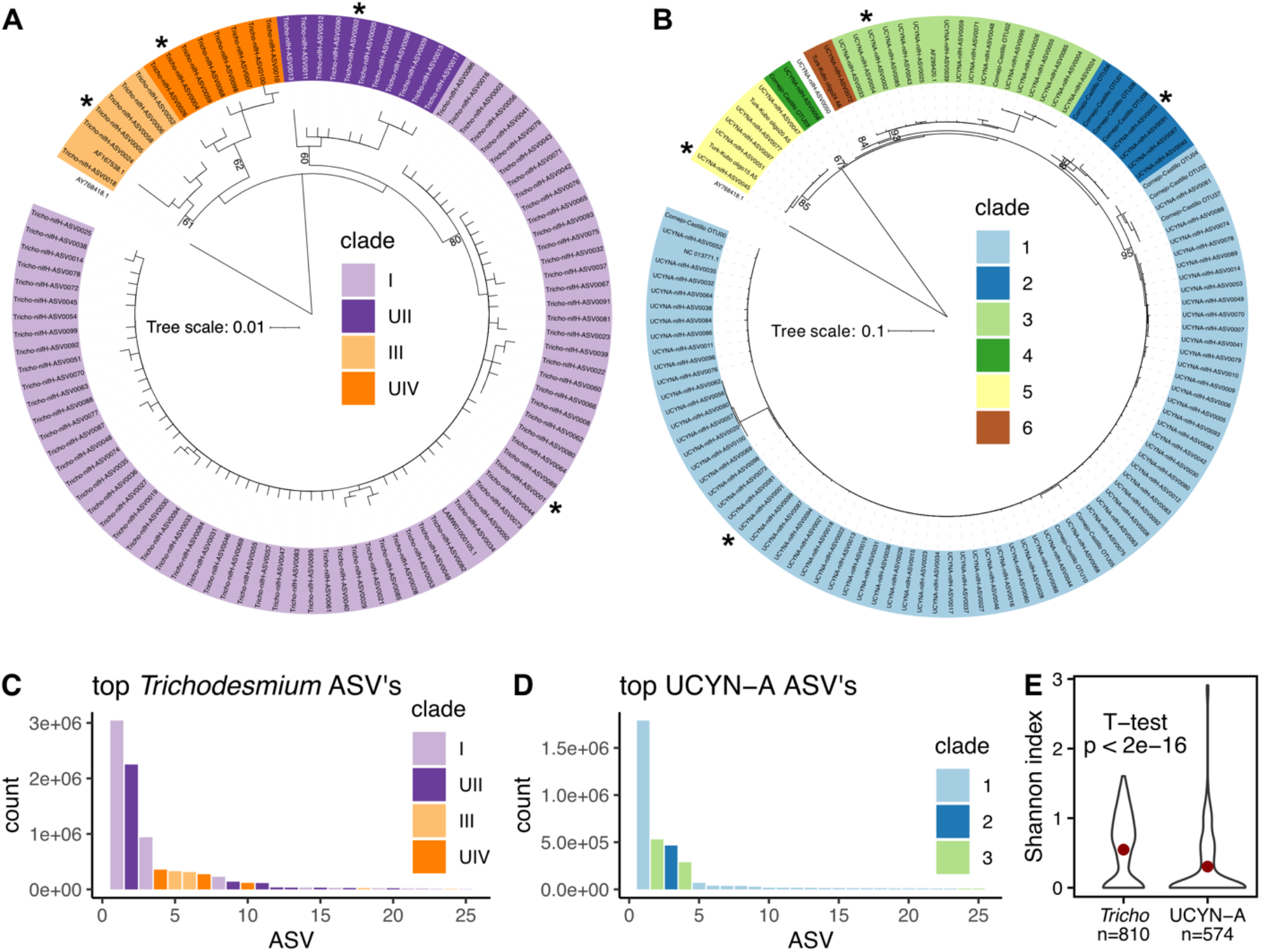
Phylogenetic tree of top 100 ASV’s in Trichodesmium (A) and UCYN-A (B). Most abundant ASV for each clade is denoted by an *. Bootstrap values out of 100 are displayed at each node. ASV naming scheme is from most abundant to least abundant. Environmental counts of top 25 ASV in Trichodesmium (C) and UCYN-A (D). Shannon diversity of *Trichodesmium* and UCYN-A (E). Mean value shown by red dot (*Trichodesmium* mean = 0.55, UCYN-A mean = 0.30).

The phylogeographies of *Trichodesmium* and UCYN-A reveal distinct regional differentiation between the different species and clades. When comparing the relative abundance of *Trichodesmium* versus UCYN-A, we see that *Trichodesmium* dominates in the northern Indian Ocean while UCYN-A is more abundant across a majority of the Pacific Ocean (Figure 3A). Across the Atlantic Ocean there is more variation between the two cyanobacteria. The AMT28 and C13.5 cruises show opposite patterns of dominance suggesting that there may be variation between *Trichodesmium* and UCYN-A due to seasonality or longitudinal gradients (Figure 1, 3A). *Trichodesmium* I was abundant in warm regions with seasonally shallow nutricline depths including the Indian Ocean and north of the equator in the Pacific and Atlantic Oceans (Figure 3B). In contrast, the opposite pattern was observed in *Trichodesmium* UII, which was abundant in the southeastern Indian Ocean near Australia and in sub-tropical to sub-arctic regions throughout the Atlantic and Pacific Oceans (Figure 3C). Clades I and II combined made up a majority of all *Trichodesmium* sequences (81.74% of total Tricho sequences)(Figure 3B, C). On the other hand, *Trichodesmium* III was abundant in the northern hemisphere of the Pacific and Atlantic Oceans, whereas *Trichodesmium* IV was abundant in the southern hemisphere (Figure 3D, E). UCYN-A1 dominated throughout the globe (Figure 3F). UCYN-A1 was the most abundant clade in samples where UCYN-A was more abundant than *Trichodesmium* (Figure 3A,F). This clade was also dominated by a single ASV (UCYNA-nifH-ASV-0001)(76% of UCYN-A1 ASVs) resulting in a fairly homogeneous UCYN-A population across the globe. UCYN-A clades 2 and 3 were primarily found in the East and West of the Indian Ocean respectively (Figure 3G, H). Each of the clades examined demonstrated a unique, regionally partitioned biogeography.

**Figure 3:**
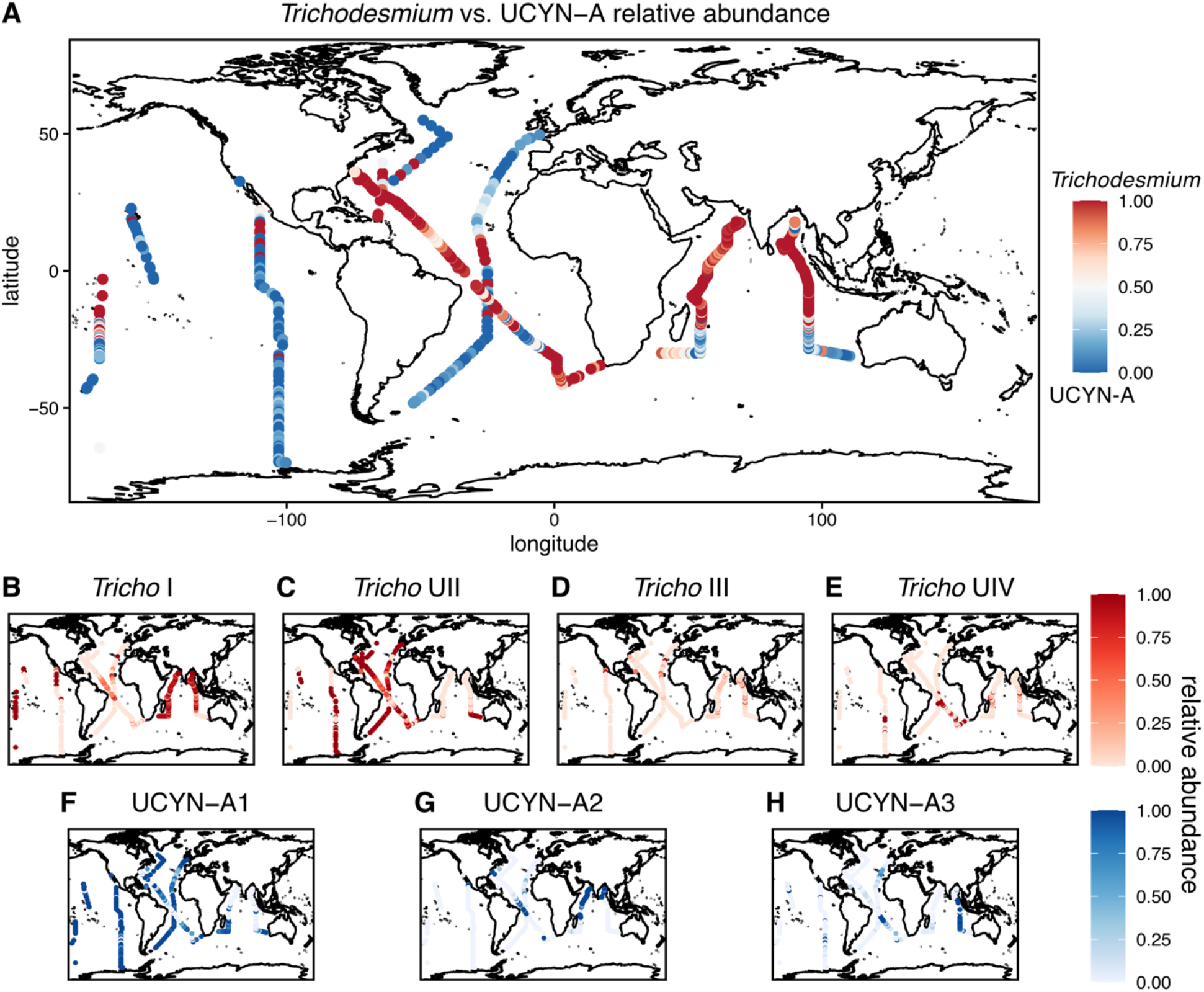
Biogeography of *Trichodesmium* and UCYN-A clades. Relative abundance of *Trichodesmium* vs. UCYNA-A (A). Relative abundance of individual clades out of total *Trichodesmium* abundance (B-E). Relative abundance of individual clades out of total UCYN-A abundance (F-H).

A comparison to environmental measurements revealed the factors shaping the diversity of *Trichodesmium* and UCYN-A. A PCoA analysis of *Trichodesmium* community diversity revealed clear separation between ocean basins with the Atlantic Ocean (AMT28, BVAL46, C13) and the Indian Ocean (I07, I09) separating across the first two dimensions (Figure 4A). We correlated environmental measurements to the PCoA results and found significant correlations for all factors compared to *Trichodesmium* (P < 0.0009). Sea surface temperature had the strongest correlation to *Trichodesmium* diversity (R^2^ = 0.32) followed by phosphorus stress (R^2^ = 0.19)(Figure 4A, Table S1). UCYN-A diversity did not partition based on ocean basin (Figure 4B, Table S2). All environmental factors had a significant correlation with UCYN-A diversity (P < 0.0009) except for phosphorus stress.

**Figure 4:**
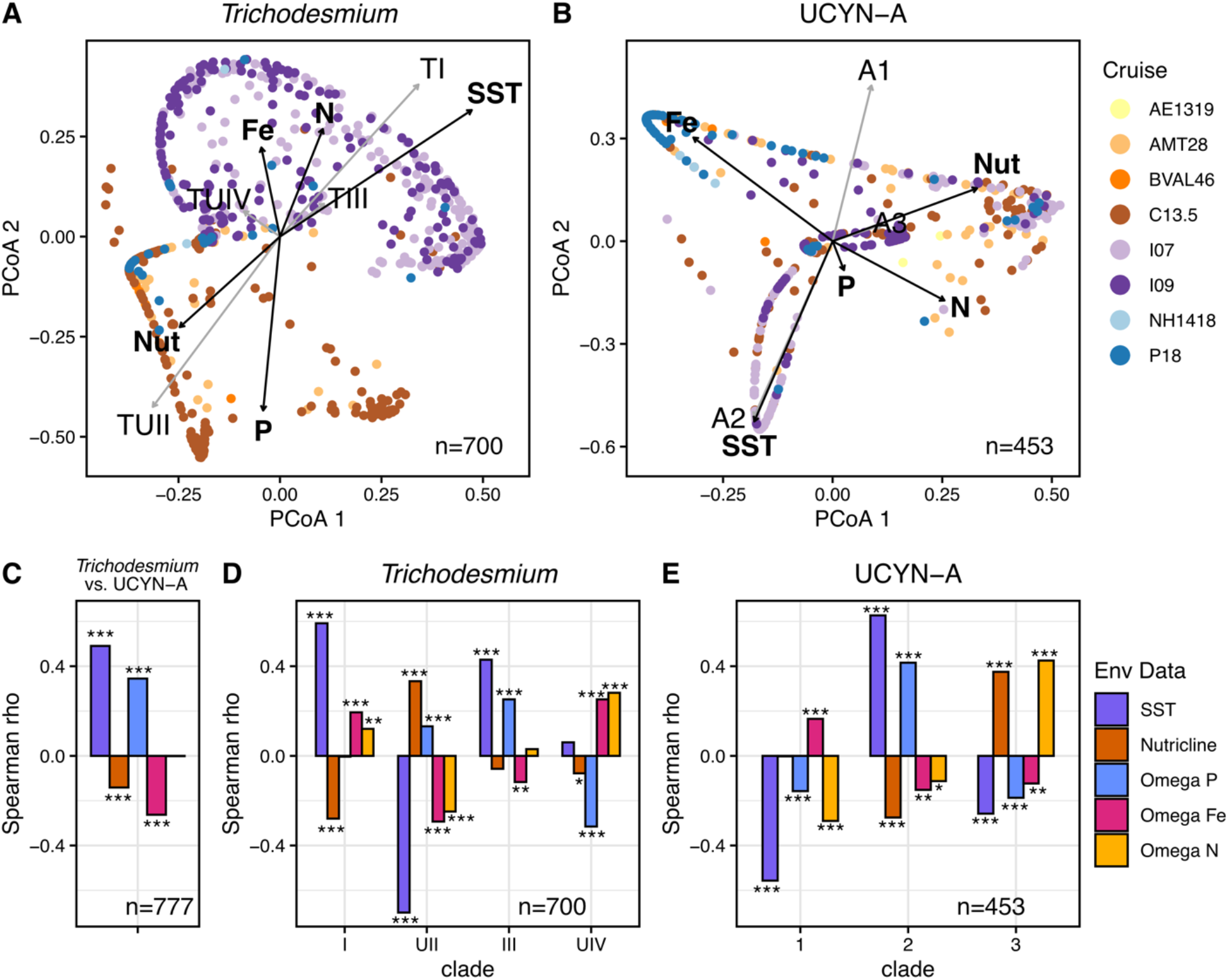
PCOA of *Trichodesmium* and UCYN-A diversity (A, B). Correlations between environmental measurements (black arrows) and clade relative abundance (grey arrows) and PCoA components, exact values can be found in (Table S1, S2)(A,B). Spearman correlations on the relative abundance of *Trichodesmium* and UCYN-A (C), *Trichodesmium* clade relative abundance (D), and UCYN-A clade relative abundance (E). Correlations are made with the following environmental data: sea surface temperature (SST), nutricline depth (nutricline), phosphorus stress (Omega P), iron stress (Omega Fe), and nitrogen stress (Omega N).

Temperature (R^2^=0.31), iron stress (R^2^=0.19), and nutricline depth (R^2^=0.13) explained the most variance (Figure 4B, Table S2). The samples are clearly partitioned across these three factors in the PCoA and show overlap between the Atlantic and Indian Ocean unlike *Trichodesmium*. A comparison of environmental factors to diversity revealed that *Trichodesmium* and UCYN-A diversity are both driven by different conditions.

We next compared the environmental drivers of *Trichodesmium* and UCYN-A. For *Trichodesmium*, we observed a significant positive correlation of sea surface temperature and phosphorus stress. In contrast, UCYN-A aligned with a deep nutricline and iron stress for (Figure 4C, Table S3). The two most abundant *Trichodesmium* clades I and UII had opposite associations with environmental factors (Figure 4D, Table S3). Clade I was more common under high temperatures, shallow nutriclines, iron stress, and nitrogen stress while clade UII was associated with lower temperatures, deeper nutriclines, and phosphorus stress (Figure 4D, Table S3). Clade III was associated with high temperatures and phosphorus stress (Figure 4D, Table S3). Clade 4 was mainly driven by nutrient conditions with a positive association with iron and nitrogen stress and a negative association to phosphorus (Figure 4D, Table S3). The globally dominant UCYN-A1 clade had a significant negative association with sea surface temperature as well as significant correlations with all nutrient stress indicators (Figure 4E, Table S3). UCYN-A2 and A3 are found in opposite parts of the Indian Ocean, which is reflected in their environmental drivers (Figure 3G, H). Clade A2 is positively correlated to high temperatures and phosphorus stress found in the northern Indian Ocean with A3 correlating to shallow nutriclines and nitrogen limitation (Figure 4E, Table S3). Thus, different environmental factors are driving the global distributions and clades of both *Trichodesmium* and UCYN-A.

## Discussion

When comparing the distributions and diversity of *Trichodesmium* and UCYN-A, we observed greater levels of microdiversity in *Trichodesmium* than in UCYN-A. This is likely due to differences in ecological strategy. *Trichodesmium* is a free-living cyanobacteria and is directly impacted by variable environmental conditions resulting in different regional selection. On the other hand, UCYN-A is an obligate symbiont with small eukaryotic phytoplankton giving it more consistent environmental conditions (Coale et al. 2024; Thompson et al. 2012). The limited diversity and dominance of a single UCYN-A ASV suggests it experiences stronger selective pressures, supporting the hypothesis that it is becoming integrated as an organelle (Coale et al. 2024). Additionally, overall *Trichodesmium* diversity is separated by ocean basin (figure 3A) while UCYN-A populations have more overlap between regions. We observed a significant correlation of *Trichodesmium* versus UCYN-A distributions separated by Fe and P nutrient stress. We may see this trade off because of the high iron requirement of the nitrogenase enzyme (Kustka et al. 2003; Raven 1988). The symbiont host of UCYN-A could be supplementing it with additional Fe allowing it to flourish in regions where *Trichodesmium* is less successful. The hypothesis that UCYN-A’s host might supplement iron aligns with existing evidence of nutrient exchanges in marine symbiotic relationships, where hosts often provide essential nutrients to their symbionts. Such interactions are well-documented in various symbiotic systems (Day and Smith 2021; Thompson et al. 2012).

While *Trichodesmium* distributions have been studied extensively at the genus-level, the biogeography of *Trichodesmium* clades has been limited (Davis and McGillicuddy 2006; Dugdale, Menzel, and Ryther 1961; Pierella Karlusich et al. 2021; Subramaniam et al. 2001). Four main clades of *Trichodesmium* have been characterized primarily using the *hetR* and ITS genes (Hynes et al. 2012; Orcutt et al. 2002). We identified four distinct clades based on the *nifH* gene (Figure 2A). Two of these clades have not been previously identified using *nifH*, and thus we inferred which clade they were based on their position on the tree relative to known clades. In particular, *Trichodesmium* IMS101 is a common laboratory isolate used in experimental work from Clade III. Here, we showed that Clades I and II were the most globally abundant while Clade III was the least abundant out of the 4 clades. The dominance of Clade I has been shown regionally, but the global distribution of Clade II has never been characterized (Gradoville et al. 2014, 2017; Hmelo, Van Mooy, and Mincer 2012; Rouco et al. 2014; Rouco, Haley, and Dyhrman 2016). This result further highlights that the use of isolate studies based IMS101 may misinform modeling studies. Our work suggests that the understudied Clade II may warrant further physiological characterization as it is abundant across much of the globe.

The observed distributions of UCYN-A clades corroborated previous regional assessments. In particular, it has been shown that a single oligotype dominates the clade UCYN-A1 (Turk-Kubo et al. 2017). We showed that this single ASV is abundant across regions where UCYN-A outcompetes *Trichodesmium*, with a link to Fe limitation (Figure 3A, 3F, 4E). The UCYN-A2 clade was initially proposed as a coastally adapted strain but has been observed in the Arctic Ocean and other high latitude waters (Cabello et al. 2016; Harding et al. 2018; Messer et al. 2015; Thompson et al. 2012). We saw a slight association between costal proximity but did not capture this clade in our high latitude samples in the Southern Ocean (Figure 3G). As in clade UCYN-A1, we observed a single dominant ASV in UCYN-A2 populations (Figure 2D). UCYN-A3 was the only clade that switched between two main ASV’s (Figure 2D). This clade has been shown to be globally distributed, but at low relative abundance compared to UCYN-A1 (Turk-Kubo et al. 2017). We found UCYN-A3 is the dominant clade in the eastern Indian Ocean. The distribution of UCYN-A3, along with a significant correlation to a deep nutricline and high N stress, suggests it has a distinct niche amongst UCYN-A populations and is not merely maintained as a low abundance subpopulation. While there are other clades in UCYN-A, we found most of the globe was dominated by UCYN-A1, UCYN-A2, and UCYN-A3 which is consistent with previous assessments (Turk-Kubo et al. 2017).

While *nifH* has classically been used to capture diazotroph diversity, there are some important caveats to consider when interpreting the results of this study. First, as with all amplicon-based sequencing, there is an unknown level of primer bias, potentially increasing the abundance of some ASVs while underrepresenting others (Angel et al. 2018). Second, the *nifH* gene is not a single copy gene and can be found in variable copy number depending on the organism (Mise et al. 2021). Additionally, *nifH* has prevalent pseudogenes and homologues in organisms that are not diazotrophs, making it an imperfect indicator of diazotroph presence (Mise et al. 2021). Nevertheless, cyanobacteria possess a particularly strong link between *nifH* and diazotroph function. Specifically, non-diazotrophic *Trichodesmium* lack the *nifH* gene, meaning any *nifH Trichodesmium* sequences are representative of the diazotrophic species (Delmont 2021). While imperfect, *nifH* is clearly linked to diazotrophic function in both UCYN-A and *Trichodesmium*, and the historical prevalence and abundance of *nifH* amplicon data provides a wealth of contextual data for amplicon-based studies.

This study provides a comprehensive analysis of the global diversity and distribution of the diazotrophic cyanobacteria *Trichodesmium* and UCYN-A, revealing intricate patterns of phylogenetic and ecological differentiation. By leveraging high-throughput sequencing of the *nifH* gene across extensive oceanic samples, we have uncovered significant clade-specific responses to environmental variables such as temperature, nutrient availability, and nutrient stress. Our findings highlight differences in biogeography and putative ecological strategies between these two diazotrophs. The dominant presence of specific *Trichodesmium* and UCYN-A clades in distinct oceanic regions underscores their ecological specialization and potential environmental role. The data presented here serve as a valuable resource for future studies aimed at understanding the functional implications of genetic diversity in marine cyanobacteria and provide a foundation for exploring their roles in biogeochemical processes under a changing climate. This work not only enhances our understanding of marine microbial biogeography but also emphasizes the need for continued research into the adaptive mechanisms that enable these cyanobacteria to thrive in diverse marine environments.

## Supporting information

Supplemental Table 1: Trichodesmium ASV counts

Supplemental Table 2: UCYN-A ASV counts

Supplemental Table 3: sample metadata

Supplemental Table 4: ASV metadata

## Acknowledgements

We thank the chief scientists L. Barbaro, B. Carter, M. Lomas, R. Sonnerup, G. Tarran, and D. Volkov and the coordinators of GO-SHIP (L. Tally and G. Johnson) and the Atlantic Meridional Transect (AMT; A. Rees) for supporting the collection of the sequencing data.

## Author contributions

A.N., L.J.U., A.A.L., and A.C.M. designed the study and wrote the manuscript. A.N. and L.J.U. analyzed the data. L.J.U. and A.C.M. supervised the study.

## Data and Code availability

All code used in this analysis can be found at https://zenodo.org/doi/10.5281/zenodo.12636629.

## Supplements

**Table S1:** Correlation between *Trichodesmium* PCOA, environmental measurements, and clade relative abundances.

| Measurement | PCoA 1 | PCoA 2 | r2 | Pr(>r) |
| --- | --- | --- | --- | --- |
| Temp_SST | 0.83107 | 0.55616 | 0.3233 | 0.000999 |
| Nutricline_1uM_Interp | -0.73753 | -0.67531 | 0.1122 | 0.000999 |
| Omega_P | -0.09499 | -0.99548 | 0.1905 | 0.000999 |
| Omega_Fe | -0.20169 | 0.97945 | 0.0517 | 0.000999 |
| Omega_N | 0.36536 | 0.93087 | 0.0835 | 0.000999 |
| Tricho_I relative abundance | 0.52505 | 0.58426 | 0.6170 | 0.000999 |
| Tricho_UII relative abundance | -0.48226 | -0.65458 | 0.6610 | 0.000999 |
| Tricho_III relative abundance | 0.16712 | 0.12605 | 0.0438 | 0.000999 |
| Tricho_UIV relative abundance | -0.13724 | 0.09741 | 0.0283 | 0.000999 |

**Table S2:** Correlation between UCYN-A PCOA, environmental measurements, and clade relative abundances.

| Measurement | PCoA 1 | PCoA 2 | r2 | Pr(>r) |
| --- | --- | --- | --- | --- |
| Temp_SST | -0.17883 | -0.52650 | 0.3091 | 0.000999 |
| Nutricline_1uM_Interp | 0.32949 | 0.15452 | 0.1324 | 0.000999 |
| Omega_P | 0.02582 | -0.08464 | 0.0078 | 0.169830 |
| Omega_Fe | -0.31770 | 0.30286 | 0.1927 | 0.000999 |
| Omega_N | 0.25505 | -0.17318 | 0.0950 | 0.000999 |
| UCYN-A_1 relative abundance | 0.14761 | 0.75838 | 0.5969 | 0.000999 |
| UCYN-A_2 relative abundance | -0.29927 | -0.85533 | 0.8212 | 0.000999 |
| UCYN-A_3 relative abundance | 0.18703 | 0.07873 | 0.0412 | 0.000999 |

**Table S3:** Spearman correlation between relative abundance and environmental measurements. For Tricho vs. UCYN-A a positive correlation means a correlation with *Trichodesmium* while a negative correlation is for UCYN-A.

| Data | Metadata | rho | pval | sig |
| --- | --- | --- | --- | --- |
| Tricho vs. UCYN-A | SST | 0.49042779 | 2.92E-48 | *** |
| Tricho vs. UCYN-A | Nutricline | -0.1411621 | 7.87E-05 | *** |
| Tricho vs. UCYN-A | Omega P | 0.34485753 | 4.05E-23 | *** |
| Tricho vs. UCYN-A | Omega Fe | -0.2629234 | 9.44E-14 | *** |
| Tricho vs. UCYN-A | Omega N | -0.0010668 | 9.76E-01 |  |
| Trichodesmium I | SST | 0.59164441 | 2.53E-67 | *** |
| Trichodesmium I | Nutricline | -0.2799336 | 4.54E-14 | *** |
| Trichodesmium I | Omega P | -0.0035073 | 9.26E-01 |  |
| Trichodesmium I | Omega Fe | 0.19408119 | 2.28E-07 | *** |
| Trichodesmium I | Omega N | 0.12052034 | 1.40E-03 | ** |
| Trichodesmium II | SST | -0.6992146 | 8.00E-104 | *** |
| Trichodesmium II | Nutricline | 0.33318219 | 1.31E-19 | *** |
| Trichodesmium II | Omega P | 0.13168934 | 4.77E-04 | *** |
| Trichodesmium II | Omega Fe | -0.2932371 | 2.39E-15 | *** |
| Trichodesmium II | Omega N | -0.2483922 | 2.65E-11 | *** |
| Trichodesmium III | SST | 0.4290588 | 1.02E-32 | *** |
| Trichodesmium III | Nutricline | -0.0578088 | 1.27E-01 |  |
| Trichodesmium III | Omega P | 0.25188706 | 1.36E-11 | *** |
| Trichodesmium III | Omega Fe | -0.1167168 | 1.98E-03 | ** |
| Trichodesmium III | Omega N | 0.03001828 | 4.28E-01 |  |
| Trichodesmium IV | SST | 0.06018263 | 1.12E-01 |  |
| Trichodesmium IV | Nutricline | -0.0777785 | 3.97E-02 | * |
| Trichodesmium IV | Omega P | -0.3153837 | 1.25E-17 | *** |
| Trichodesmium IV | Omega Fe | 0.25186672 | 1.37E-11 | *** |
| Trichodesmium IV | Omega N | 0.28142719 | 3.29E-14 | *** |
| UCYN-A1 | SST | -0.5566254 | 3.25E-38 | *** |
| UCYN-A1 | Nutricline | -0.0021103 | 9.64E-01 |  |
| UCYN-A1 | Omega P | -0.1572629 | 7.83E-04 | *** |
| UCYN-A1 | Omega Fe | 0.16498373 | 4.22E-04 | *** |
| UCYN-A1 | Omega N | -0.2902482 | 3.04E-10 | *** |
| UCYN-A2 | SST | 0.62666607 | 8.55E-51 | *** |
| UCYN-A2 | Nutricline | -0.275072 | 2.62E-09 | *** |
| UCYN-A2 | Omega P | 0.41565014 | 2.38E-20 | *** |
| UCYN-A2 | Omega Fe | -0.1516123 | 1.21E-03 | ** |
| UCYN-A2 | Omega N | -0.112704 | 1.64E-02 | * |
| UCYN-A3 | SST | -0.2578374 | 2.59E-08 | *** |
| UCYN-A3 | Nutricline | 0.37495422 | 1.44E-16 | *** |
| UCYN-A3 | Omega P | -0.1870853 | 6.17E-05 | *** |
| UCYN-A3 | Omega Fe | -0.1233941 | 8.56E-03 | ** |
| UCYN-A3 | Omega N | 0.42529494 | 2.52E-21 | *** |

